# Gasdermin D deficiency attenuates development of ascending aortic dissections in a novel mouse model

**DOI:** 10.1101/2024.08.22.609270

**Authors:** Muhammad J. Javed, Rachael M. Howard, Hua Li, Laura Carrasco, Marvin L.S Dirain, Gang Su, Guoshuai Cai, Gilbert R. Upchurch, Zhihua Jiang

**Affiliations:** Division of Vascular Surgery and Endovascular Therapy, University of Florida College of Medicine, Gainesville, FL 32601, United States; Division of Thoracic and Cardiovascular Surgery, University of Florida College of Medicine, Gainesville, FL 32601, United States; Malcom Randll VA Medical Center, Gainesville, FL 32608

**Keywords:** aortic dissection, cell death, neutrophil, immune response, model

## Abstract

**Background:** Thoracic aortic dissection (TAD) is a silent killer. Approximately two-thirds of the cases occur in the ascending aorta (i.e. type A dissection) and majority of them are unrelated to genetic mutations. However, animal models of spontaneous type A dissection are not widely available. In the present study, a novel mouse TAD model was created. Further, the role of gasdermin D (GSDMD) in TAD development was evaluated.

**Methods:** TADs were created by treating ascending aorta of adult mice (C57BL/6J) with active elastase (40.0 U/ml) and β-aminopropionitrile (Act E+BAPN). The temporal progress of the TAD pathology was rigorously characterized by histological evaluation and scanning electron microscopy, while potential mechanisms explored with bulk RNA sequencing of specimens collected at multiple timepoints. With this novel TAD model, further experiments were performed with Gsdmd^−/−^ mice to evaluate its impact on TAD formation.

**Results:** The ascending aorta challenged with Act E+BAPN developed pathology characterized by an early onset of intimomedial tears (complete penetration) and intramural hematoma, followed by progressive medial loss and aortic dilation. Ingenuity Pathway Analysis and functional annotation of differentially expressed genes suggested that a unique inflammatory micro-environment, rather than general inflammation, promoted the onset of TADs by specifically recruiting neutrophils to the aortic wall, while the pathology at the advanced stage was driven by T-cell mediated immune injury. Gsdmd^−/−^ attenuated medial loss, adventitial fibrosis, and dilation of TADs. This protective effect was associated with a reduced number of TUNEL (terminal deoxynucleotidyl transferase dUTP nick end labeling) positive cells and T-cells in TADs.

**Conclusions:** A novel mouse TAD model was created in the ascending aorta. It produces a unique microenvironment to activate different immune cell subsets, promoting onset and subsequent remodeling of TADs. Consistently, Gsdmd^−/−^ attenuates TAD development, with modulation of cell death and T-cell response likely acting as the underlying mechanism.

**Graphical Abstract:** 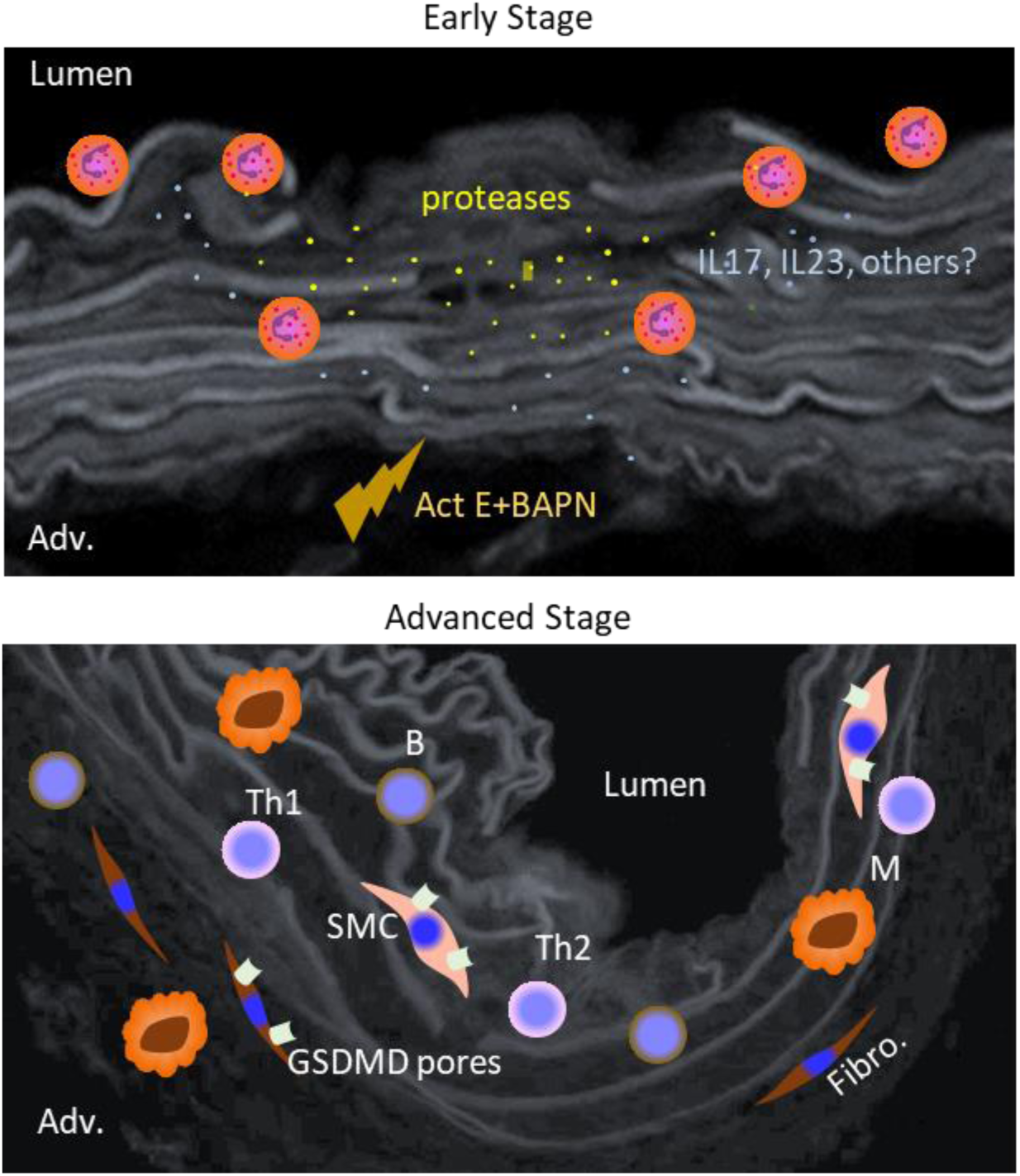

## Introduction

Aortic dissection is a silent killer, with an estimated incidence of 30 to 50 cases per 1 million person-years^1^. The thoracic aorta, particularly the ascending aorta is most frequently affected^2^. Although it is a relatively uncommon cardiovascular disease, it represents one of the major catastrophic life-threatening events. Currently, diagnosis of aortic dissections can only be made after their occurrence due to the lack of biomarkers capable of identifying the vulnerable population. Despite the prompt response of emergent medical care, approximately, 50% of the cases die before reaching a hospital^3^. Improvement of the clinical outcome of aortic dissections remains an unmet need.

Our current understanding of the etiology of aortic dissections is quite elusive. While roughly 25% of the cases may be attributable to genetic mutations, the cause for majority of the cases is unknown^4^. This lack of knowledge poses significant challenges in creating animal models capable of faithfully recapitulating the pathophysiology of human aortic dissections. Multiple animal models have been developed for studying thoracic aortic dissections (TADs). Among them, the angiotensin II (AngII) model is the most widely adopted, with modifications to the initial design being made to increase the rate of TAD formation^5,6^. Because of the limitations of individual models, we and others developed an alternative model by inducible deletion of SMC (smooth muscle cell) specific TGF-β receptors^7–9^, for cross-model validation of the scientific findings. However, this approach is time-consuming. Off-target genome modifications may occur in the genetically modified animals and confound data interpretation^10^. These shortcomings have suggested a need for alternative options.

TAD is a separate aortic disease that differs from the thoracic aortic aneurysm (TAA) in etiology, pathophysiology, and clinical management^4,11^. However, some deleterious cell events are shared by the two distinct pathologies, driving medial degeneration and rendering the aortic wall prone to dissection and/or dilation. For example, cell death driven by various programs has been implicated as a significant contributor to the development of TADs and TAAs^12^. Gasdermin D (GSDMD), a membrane pour-forming molecule when activated, is an executioner protein for pyroptosis^13^. Recent studies suggest that GSDMD may be activated by pathways other than pyroptosis, for instance, panoptosomes^14^. Other studies have documented GSDMD as a driver for AAA formation^15,16^ and rupture^17^, but its role in the development of TADs has not been explored. In the present study, we developed a novel mouse TAD model to study the role that GSDMD plays in TAD formation.

## Method

### Data Availability

Data supporting the findings of this study are available from the corresponding author upon reasonable request.

### Animal Studies

This study conforms to the Guide for the Care and Use of Laboratory Animals of the National Institutes of Health. The Institutional Animal Care and Use Committee of the University of Florida approved all procedures. Mice used in this study were on a C57BL/6J background. Both wild type (WT) and *Gsdmd^−/−^* mice were purchased from Jackson laboratory and were allowed to acclimate for at least one week prior to enrollment in the study.

Elastase, active (Act E) or heat inactivated (Inact E) was delivered to the outer surface of the ascending aorta via minimally invasive surgeries under a dissecting microscope. Briefly, mice were anesthetized by continuous inhalation of 1.5-2.0% isoflurane and secured on a supine position. After removing the fur and disinfection of the surgical area, a midline incision (around 1.0-1.5 cm) was made, extending from the lower neck to the upper chest. A straight scissor was used to cut the manubrium longitudinally, followed by opening the manubrium with a custom-made retractor. Thymus was separated bluntly from the anterior wall to expose the ascending aorta and aortic arch, followed by isolating them from peri-aortic tissue with fine forceps. A 30.0 μl elastase was then loaded to two cotton patches, with each being placed to wrap the back and front of the ascending aorta. After a five-minute incubation, the cotton patches were removed. A 4-0 absorbable suture was used to restore the manubrium and to close the skin incision. Ethiqa-XR, a slow-releasing version of buprenorphine, was administered via a s.c. injection to alleviate pain and stress. In our hands, the successful rate is around 90%, with no surgery-related post-op death noted so far. Beta-aminopropionitrile (BAPN, 0.2%) was delivered on the day of surgery via drinking water and refreshed twice weekly.

### Ultrasound imaging

Animals were anesthetized via inhalation of 1.5-2.0% isoflurane and placed in a supine position on a heated platform. Naire was used to remove the fur from the area of imaging and then, wiped out with warm water. A high-resolution Vevo 2100 Imaging System with a MS550D (central frequency: 40 MHz) linear array transducer (VisualSonics) was then utilized to scan aortas under B-mode. Ascending aortas were imaged via the right parasternal long axis view, with the transducer finely adjusted to a position that captures the largest possible width of the ascending aorta. A cine-loop was then recorded and reviewed to select the frame that displays the widest lumen. On the selected image, lumen diameter was measured using the straight-line tool, with the line placed and oriented so that the length of the line equals the largest lumen diameter of the ascending aorta^8,9^.

### Histology

Evans blue staining was performed as described previously^5^. Briefly, Evans blue solution (5%) was administered through tail vein injection 30 minutes before tissue collection. During tissue collection, saline was injected through the left ventricle to thoroughly remove Evans blue from the circulatory system, followed by perfusion fixation with 10% neutral buffered formalin. The TADs were cut open longitudinally and spread on slides with the luminal side facing up. After mounted with coverslips, the specimens were evaluated microscopically (Zen lite 2012, Carl Zeiss, Oberkochen, Germany) for Evans blue extravasation and presence of intimal/medial tears.

Van Gieson and Prussian blue staining were performed on paraffin sections. While a standard protocol was followed for Va Gieson staining, a kit (ab150674, Abcam, Cambridge, United Kingdom) was used for Prussian blue staining by following the manufacture’s instruction.

### Morphometry

Cross-sections (5.0 µm) were collected at locations 0, 100, and 200 µm from the proximal end of the TADs, with the point displaying the first complete circle taken as the position-0 (p0). Specimens were stained using the Van Gieson protocol and imaged with a Zen blue software (Carl Zeiss, Oberkochen, Germany). Both internal and external elastic laminae were traced to obtain area of media and length of the external elastic laminae (EEL), while area of the adventitia was obtained by tracing outer boarder of the layer with dense matrix. For each TAD sample, the area of the media and adventitia was normalized to the length of EEL. To evaluate medial loss, surface length was measured for areas where the media contains ≤ 4 layers of elastic lamina (the mouse ascending aorta generally has 8 layers of elastic laminae) or displays ≥ 50% reduction in thickness. This criterion was set so that the starting and the ending points are standardized when lines were drawn to measure medial loss. For a given section, length of lines was summed up, followed by normalization to EEL.

### TUNEL Assays

TUNEL assays were performed with a One-step TUNEL In Situ Apoptosis Kit (E-CK-A322, Elabscience) following manufacturer’s instructions. Staining was evaluated by two observers who were blinded of the treatment information and counted number of TUNEL^+^ cells on individual sections. An average of the counts obtained from the three locations was calculated for statistical analysis.

### Immunohistochemistry (IHC) and immunofluorescence (IF) Assays

IHC and IF assays were performed on paraffin sections collected at the location of P100, following protocols established in our laboratory previously^9,18^. Briefly, specimens were rehydrated and antigens unmasked by incubating specimens in Citra buffer (pH 6.0, H3300, Vector Laboratories) heated in a pressure cooker. After blocking non-specific bindings, specimens were incubated with primary antibodies at 4°C overnight, followed by probing secondary antibodies conjugated with Alexa Fluro or biotin. For IHC assays, antigen-specific signals were amplified with an ABC kit (PK-6100, Vector Laboratories) and developed with a DAB kit (SK-4100, Vector Laboratories) Nuclear counterstain was completed with hematoxylin and DAPI (D9542, Sigma-Aldrich) for IHC and IF labeling, respectively.

IHC assays were evaluated under a bright field microscope by two observers blinded of grouping information. Both observers graded density of positive cells using a 5-scale scoring system. In cases where the difference was greater than 2 points between the two observers, assays were graded by a third reviewer. An average score was calculated to represent positivity of the assay on each specimen. IF assays were graded following the same protocol as used for IHC assays except reviewing under a fluorescent microscope.

### Bulk RNA sequencing

Total RNAs were extracted using a TRIzol plus RNA Purification kit (12183555, Invitrogen) following manufacturer’s instructions. An aliquot was collected for every sample and analyzed with an RNA 6000 Pico Kit (5067-1513, Agilent) on an Agilent 2100 Bioanalyzer (Agilent, CA). All RNA samples produced a RIN > 8.0 while those entering next step had a RIN ≥ 8.5.

Illumina Stranded mRNA Prep and Ligation Kit (Ref# 20040534; Illumina, CA) was used to prepare sample libraries following the manufacturer’s recommendations. Briefly, high-quality total RNA (100-300 ng/µL) was captured and purified, followed by reverse transcription to convert the captured mRNA into first and second strands of cDNA. The resulting double stranded cDNA molecules were then enzymatically fragmented in preparation for unique dual indexing using IDT for Illumina RNA UD Set Ligation Indexes (Ref# 20040534, Illumina). To facilitate ligation of the barcoded adapters, adenine (A) nucleotides was added to the 3ʹ ends of the blunt fragments preventing them from ligating to each other during adapter ligation. Additionally, a corresponding thymine (T) nucleotide on the 3ʹ end of the adapter provided a complementary overhang for ligating the adapter to the fragment. The ligated samples were subsequently enriched through amplification (13x PCR cycles) on Bio-Rad Thermal cycler (T100; Bio-Rad, CA) and purified using AMPure XP magnetic beads (Ref# A63881; Beckman Coulter, CA). The average fragment length of libraries was measured to be ∼300–400 bp using the DNA 1000 Kit (Agilent, CA) on Agilent 2100 Bioanalyzer. Finally, the optimized libraries were pooled to a final concentration of 750pM. Afterwards, they were loaded onto Illumina’s P2 flow cell/200-cycle reagent cartridge (Ref# 20100986, Illumina) for sequencing on the NextSeq 1000 instrument (Illumina, CA). The run setup for the Illumina NextSeq 1000 was set to a length of 100bp reads (Read1: 100; Index1: 10; Index2: 10; Read2:100). Denature and Dilute On Board was enabled and a 2% PhiX spike-in was added as positive control. To ensure quality, multi-QC Analysis was conducted for each sequence run using Illumina’s Dragen RNA software (Illumina). Once sequencing was completed, the raw sequence base call files (BCL) were converted to FASTQ files using Illumina’s bcl2fastq conversion software (Illumina, CA) as input for downstream bioinformatics analysis.

### Bioinformatics

The raw sequencing reads were quality controlled using FastQC (http://www.bioinformatics.babraham.ac.uk/projects/fastqc/) and mapped to the mouse GRCm39 genome assembly as well as Gencode M32 annotations using STAR aligner^19^. RSEM^20^ were applied to summarize gene counts, which were further normalized using the TMM method^21^ and gene-wise compared between experimental groups using edgeR^22^ based on negative binomial generalized linear models. The Benjamini & Hochberg procedure was applied to adjust P-values for multiple comparison. Differentially expressed Genes were identified with an FDR of 10 and an adjusted *P*<0.05. Principal component analysis was performed on the scaled logarithm transformed FPKM (fragments per kilobase per million) value and samples were examined in the space of the first three principal components (PC1, PC2 and PC3).

IPA (Ingenuity Pathway Analysis) “Core Analysis” was performed to identify canonical pathways, and their upstream regulators predicted to be accountable for DEGs^23,24^. A cutoff of 2.0 and 10.0 was set for Z score and P value, respectively. Pathways associated with specific diseases, such as arthritis and cancer, were excluded from the graphical summary due to the irrelevance of those themes to the present study. Functional annotation of DEGs was performed with DAVID ^23,24^. With an EASE score of 0.01 and a minimum gene count of 10, KEEG pathways were enriched and the annotation chart exported as a text file. Following collapsing of redundant themes, data were plotted as a dot plot.

## Statistical Analysis

All data are expressed as the mean ± SEM. Statistical analyses were performed using Sigma Plot 14.0 (San Jose, CA, USA). Datasets were evaluated using normality and equivalence variance testing. For those failing this evaluation, logarithmic and exponential transformations were performed to meet these requirements. Student’s t-test, one-way ANOVA, two-way repeated measures ANOVA, and Mann-Whitney Rank Sum test were performed, when appropriate, with Holm–Sidak analysis being used for post hoc tests. P<0.05 was considered statistically significant.

## Results

### Elastase and BAPN work in concert to promote TAD formation

Elastase is one of the proteases that are commonly used for the creation of animal models of TAAs and AAAs^25,26^. These models appear to develop aortic aneurysms, with rare aortic dissection or rupture occurrence. In the abdominal aorta, the combined challenge of elastase and BAPN induces AAAs with a phenotype featuring persistent aortic dilation and luminal thrombosis^27^. These results predicted that systemic delivery of BAPN in combination with local elastase insult would yield a model of TAA. However, the same treatment resulted in a phenotype of dissection rather than aneurysm dilation in the ascending aorta, and therefore, the term TAD is used to indicate the phenotype throughout the present study.

The ascending aorta was approached through a midline incision (1.0-1.5 cm) crossing the lower neck and the upper chest and isolated without opening the chest. Elastase (30.0 μl) was loaded to two cotton patches (15.0 μl/piece) and delivered to the outer surface of the ascending aorta by wrapping it from both the anterior and the posterior side for five minutes. Since both low and high concentrations of elastase were used to induce AAAs^27,28^, we performed a set of experiments to determine the appropriate dosage for TAD induction. C57BL/6J mice (male, 11-14 weeks of age) were randomized to three groups (n=3-5 per group), with the ascending aorta exposed to active elastase (Act E) at a low (30.0 μl at 4.0 U/ml) or a high dose (30.0 μl at 40.0 U/ml), or heat inactivated elastase (Inact E, 30.0 μl heated high dose solution). BAPN (0.2%) was administered to all three groups, beginning on the day of surgery. Ultrasound imaging showed that the low dose elastase failed to promote aortic dilation in a four-week follow-up period. The high dose elastase resulted in aortic dilation. However, the temporal pattern of aortic dilation differed from that documented for elastase-induced AAAs. Specifically, early dilation (< 1 week) was not dramatic. The overall expansion was much less than 100% (Figure S1A) as seen in the AAA model^28^.

Because of the different aortic phenotype, we went on to perform rigorous model characterization. C57BL/6J mice (male, 11-14 weeks of age) were randomized to receive treatments as follows (n=6-8 per group): Act E+BAPN, Act E, Inact E +BAPN, or Inact E. Ultrasound imaging showed that both the Act E+BAPN and the Act E promoted an early (in a week), moderate aortic dilation (23 ± 4 % and 23 ± 3 % for Act E+BAPN and Act E, respectively). Thereafter, aortas exposed to Act E+BAPN continued to expand and caused a 71 ± 13% increase in lumen diameter in four weeks. In contrast, those exposed only to Act E halted the progression, showing insignificant changes in diameter during the rest of the follow-up period. The groups challenged with Inact E +BAPN Inact E did not result in significant aortic dilation in the entire four weeks (Figure 1A). Peri-aortic adhesion and aortic dilation are the typical pathologies noted during gross examination. Absorbed intramural hemorrhage (yellow coloration of the aortic wall) was detected occasionally (Figure 1B and S1B). Fresh intramural hemorrhage (presence of red clot or dots) was not detected at the four-week time point. Aortic rupture was not noted for this cohort during the four-week follow-up.

**Figure 1.**
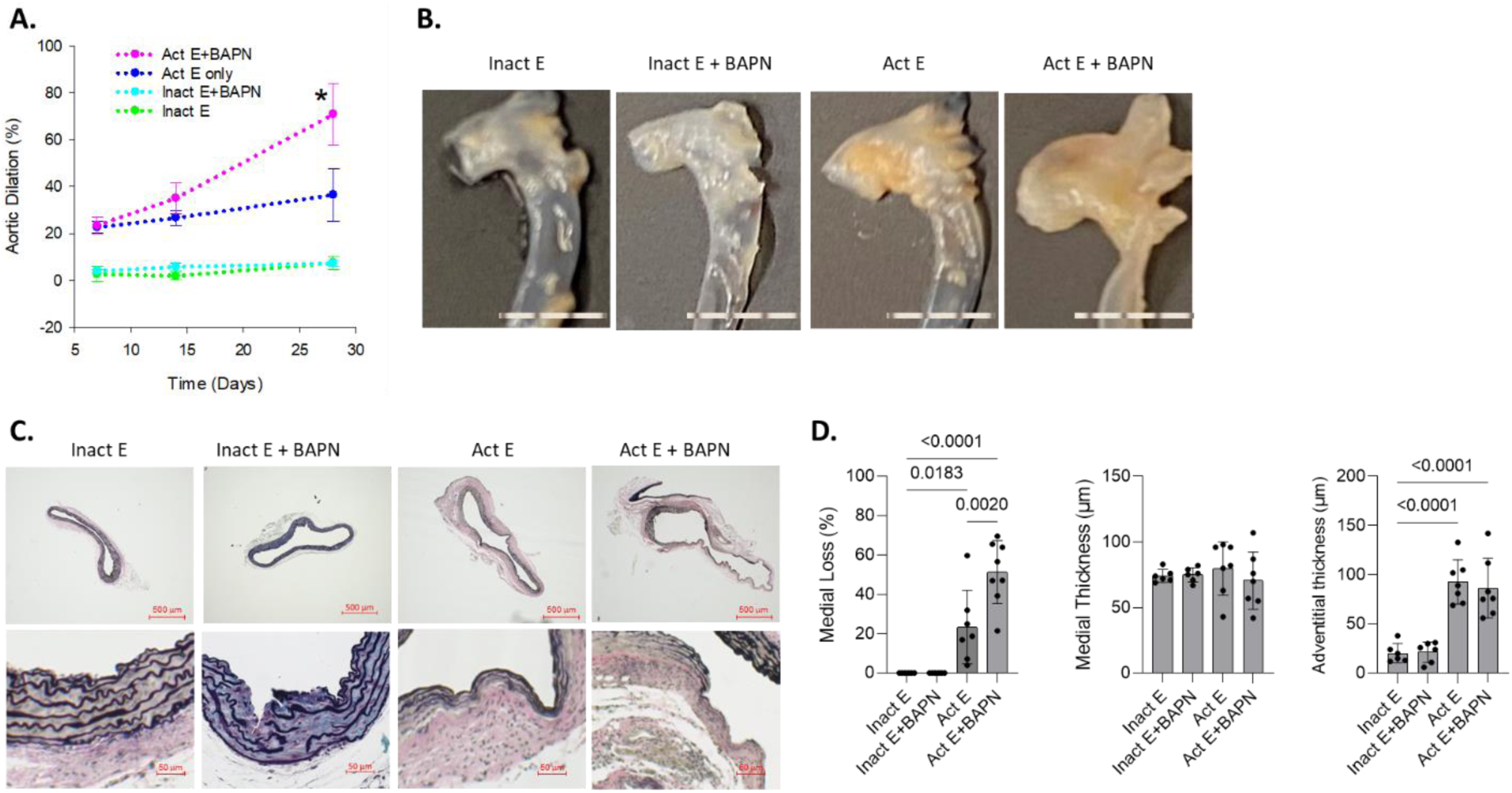
A reliable mouse TAD model created with minimally invasive surgery. Specimens were collected four weeks after the TAD induction. A, dilation of TADs challenged with indicated conditions (C57BL/6J, male, 10 to 14 weeks old, n=6-8). Data was analyzed using two-way RM ANOVA. Time: *P*<0.001; treatment: *P*<0.001; Time × treatment: *P*<0.001; \**P*=0.005, Act E+BAPN *vs*. Act E. Inact E: heat-inactivated elastase; Act E: activate elastase without BAPN; Act E+BAPN: active elastase with BAPN. B, representative gross images of aortas subjected to indicated treatments. Ruler scale: 1.0 mm. C, Van Gieson staining of TADs. High power images (lower row) depict typical pathologies developing under indicated conditions. D, morphometric evaluation. Data was analyzed using one way ANOVA.

To further characterize the TAD pathology, we performed serial sectioning at a 100 μm interval for specimens collected at the four-week time point. An example is provided in Figure S1B and S1C. Elastic fiber breaks, intimomedial tears, medial thinning or complete loss, and adventitial fibrosis were readily detectable in aortas exposed to Act E with or without BAPN, while not penetrating to those treated with Inact E only (Figure 1C). Blood clots were occasionally located between elastic laminae (Figure S1D) in TADs induced by Act E+BAPN (3 in 8) or Act E (2 in 8). This pathology was defused along the circumference of the aorta except the area adjacent to the pulmonary artery, where medial thinning or complete loss was less frequently observed. Along the long axis of TADs, the whole ascending aorta presented the pathologies described above with little variation, except the proximal end, where medial loss was not detected in some cases (Figure S1C). Aortas treated with Inact E +BAPN were not free of pathology. Elastic fiber breaks and partial intimomedial tears were frequently detected, although medial thinning was absent in these aortas (Figure 1C). Because of the massive medial loss, we decided not to count elastic fiber breaks or intimomedial tears. Rather, we quantified severity of medial loss (defined as area with ≤3 layers of elastic lamina, Figure S2A and S2B). Act E+BAPN or Act E alone resulted in a significant medial loss, with the pathology much more pronounced in TADs in the presence of BAPN, while neither Inact E nor Inact E+BAPN caused discernible medial loss. Interestingly, despite evident medial loss, the overall medial mass of TADs was not significantly different from aortas exposed Inact E or Inact E +BAPN (Figure 1D), indicating thickening of the residual media in TADs. Adventitial thickening, particularly in the area without medial loss, was evident in Act E as well as Act E+BAPN TADs (Figure 1C). Consequently, these TADs displayed much thicker adventitia than aortas exposed to Inact E or Inact E +BAPN (Figure 1D).

Inflammatory infiltrates in TADs were labeled with immunohistochemistry (IHC) assays. T-cells (CD4+ or CD8+), B-cells (CD19+), and macrophages (CD68+), were readily detectable in aortas challenged with Act E+BAPN and more frequently located in the adventitia (Figure S3A). Compared with aortas exposed to Inact E, Inact E +BAPN, or Act E, TADs caused by Act E+BAPN displayed significantly more B-cells and macrophages (Figure S3B). Neutrophils (Ly6b.2+) were not detected on any of the specimens (data not shown). Additionally, we evaluated angiogenesis with IHC assays for expression of the CD31. “Floppy vessels” were frequently found in the adventitial layer of TADs and aortas challenged with Act E, while not detected in those exposed to Inact E or Inact E+BAPN (Figure S4A). Quantitatively, TADs and aortas exposed to Act E had a greater degree of angiogenesis than aortas under other conditions (Figure S4B). Collectively, these results suggest that TADs suffer chronic inflammation at the advanced stage.

### Act E+BAPN causes early onset of intimomedial tears and aortic wall dissection

The pathology observed for advanced TADs induced by Act E+BAPN is distinct from TAAs created with the same dosage^26^, but was very similar to what we documented for advanced TADs induced by chronic AngII infusion^5^. To further characterize the aortic phenotype, we evaluated the pathology occurring at the early stage. First, we evaluated integrity of the endothelial barrier with Evans blue staining and En face microscopy as described previously^8^. TADs were collected on d3 from WT mice exposed to Act E+BAPN or Inact E +BAPN and processed for bright field or scanning electron microscopy (SEM). For the Act E+BAPN group, fresh intramural hematoma (presence of blood within the aortic wall) was noted in 2 out of the 6 specimens. Isolated blue areas with or without intimomedial tears were found in the anterior and posterior aortic wall. Intimomedial tears generally oriented with the direction of blood flow, while deferring in length (30-420 μm), depth (partial to full media), and frequency (6-11 tears/sample) among the specimens. Tears were always present in areas with fresh intramural hematoma, but most tears were not associated with this event (Figure 2A and 2B). Scanning electron microcopy confirmed intimomedial tears in TADs. Adherent leukocytes and thrombi were located on surface of the media lacking intimal coverage (Figure 2C). Elastic fiber breaks and dissemination of the aortic wall were readily detectable in cross sections of the d3 TADs under fluorescence microscopy (Figure 2D and 2E). Pathologies described above were not observed in aortas exposed to Inact E +BAPN (data not shown). These results demonstrate that Act E+BAPN causes development of TADs, instead of TAAs, in the ascending aorta.

**Figure 2.**
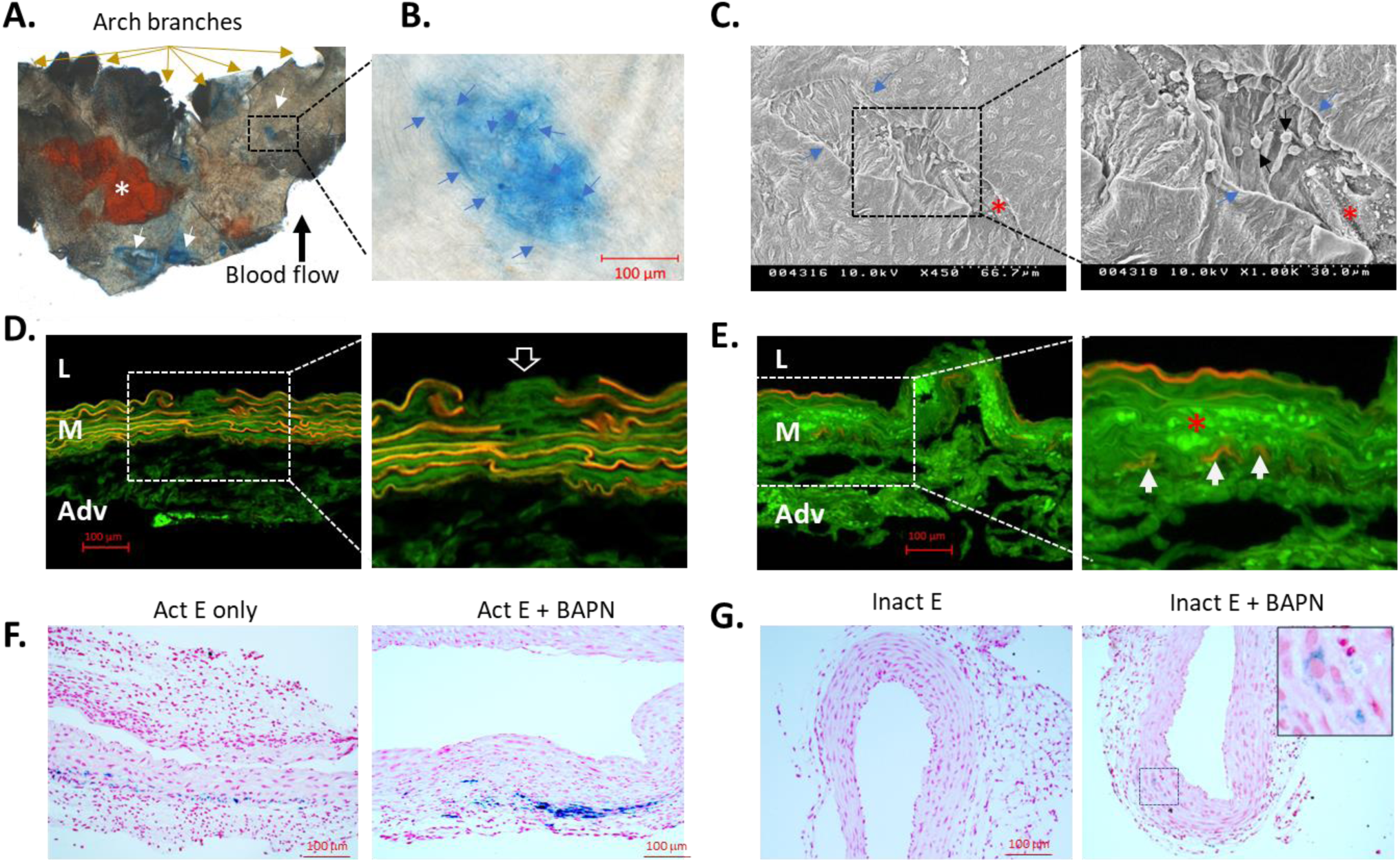
Pathology identified TADs at d3. TADs were collected from WT mice exposed to Act E+BAPN (n=6) or Inact E+BAPN (n=6). A, En face imaging of TADs (n=3/treatment). Yellow arrows indicate branches of the aortic arch. The black arrow depicts the direction of the blood flow. The white star indicates an area with fresh intramural hematoma. Areas with intimomedial tears or impaired endothelial barrier function were stained in blue due to extravasation of Evans blue (white arrows). B, a magnified view of the boxed area in A, showing an intimomedial tear highlighted by Evans blue staining. Blue arrows point to the edge of an intimomedial tear. C, Scanning electron microscopy of the luminal surface of TADs (n=3/treatment). Blue arrows indicate the edge of the intimomedial tear while the red star depicts a thrombus. A magnified view of the boxed area is shown on the right, where black arrows point to adherent white blood cells while the red star depicts a thrombus containing platelets (small white bodies) and amorphous fibrin. D and E, fluorescence imaging of d3 TADs subjected to Evans blue staining *in vivo*. Evans blue penetrated full thickness of the media, with the stained elastic laminae emitting red fluorescence. The open arrow points to an intimomedial tear (D). Blood clot (indicated with a red star (E) trapped in the medial layer and emitted bright green fluorescence. White arrows point to the layer of external elastic lamina. F and G, Prussian blue staining of d28 TADs. Blue: ferric irons deposited by dead red blood cells.

Following the creation of this TAD model, we used it in multiple studies. Gross examination was performed for all TADs that were collected at various timepoints (n=41, Table 1). Aortic enlargement and peri-aortic adhesions were observed in all TADs. An interesting finding, however, is the timing of the onset of aortic dissections. Fresh hematoma was noted in 7 out of the 9 TADs collected on d7, but rarely detected at d14 or d28. Absorbed hematoma (defined as yellow coloration of the aortic wall) was absent in the first week of surgery, but grossly evident only in some TADs collected on d14 or d28. To detect resolved hematomas, we performed Prussian staining on d28 TADs. Positive staining was observed in 4 out of 8, 4 out of 7, and 2 out of 6 TADs induced by Act E+BAPN, Act E, and Inact E +BAPN, respectively, while not detected in any of the aortas exposed to Inact E (Figure 2F and 2G), suggesting that Prussian staining could not catch all ever-existing hematomas in TADs.

**Table 1:**
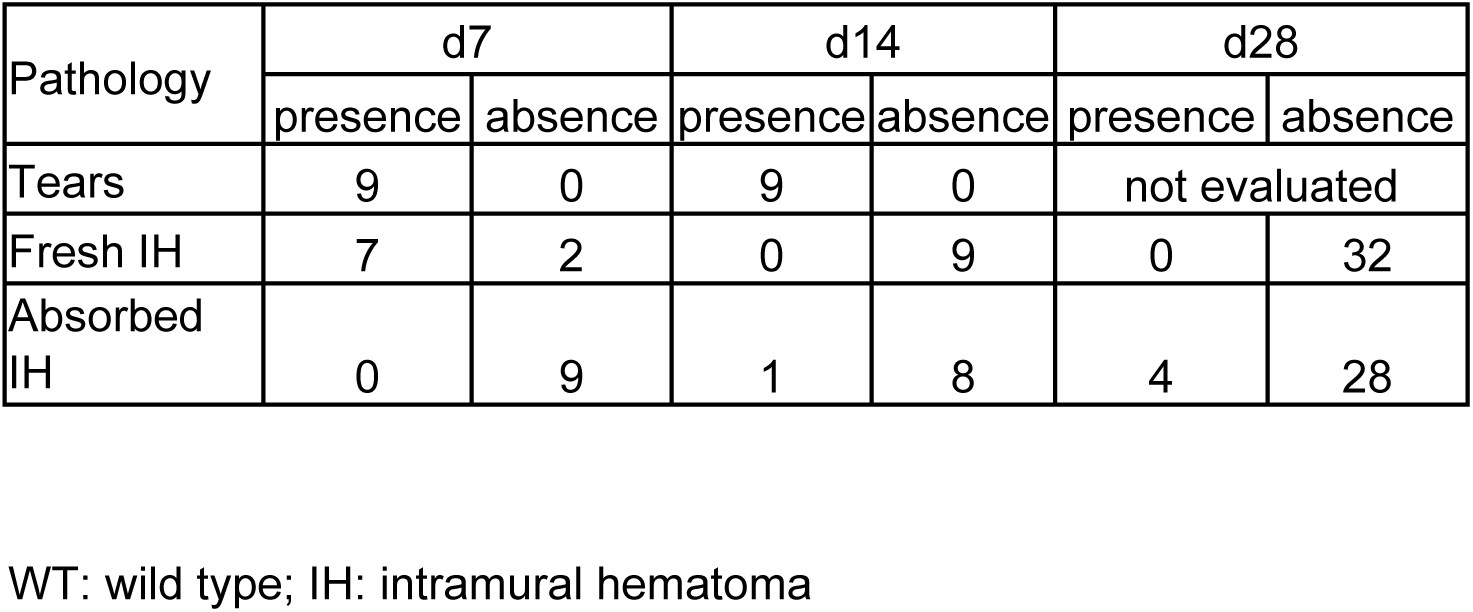
Gross pathology detected in WT TADs induced by Act E+BAPN.

### Act E+BAPN perpetuates inflammatory response in the aortic wall

An interesting finding on aortas challenged with various conditions is that Act E+BAPN and Inact E both stimulated inflammation in the aortic wall, but only Act E+BAPN induced TAD formation. To explore mechanisms responsible for the phenotypic disparity, we performed bulk RNAseq for aortas exposed to those conditions and compared their gene expression profiles. TADs were collected at d7 and d14 while naïve aortas were included as a baseline reference. Principle component analysis showed that TADs cluster together by stimulation (Figure S5), indicating distinct gene signatures among groups subjected to different treatments. Time-dependent clustering was evident for aortas exposed to Inact E, but not for TADs (Figure S5), suggesting that the overall gene expression profile does not differ between TADs collected on d7 and d14.

To explore treatment- and time-dependent regulation of pathways and events, we performed Ingenuity Pathway Analysis (IPA) with differentially expressed genes (DEGs). The first set of analyses was carried out to evaluate early (d7) regulation of gene expression by Inact E and Act E+BAPN, and then analyzed temporal changes of gene expression from d7 to d14. With an FDR of “10” and a significance level of 0.05, 9,255 and 11,106 genes were differentially expressed in aortas exposed to Inact E (Figure 3A) and Act E+BAPN (Figure 3C), respectively, at d7, compared with naïve aortas. IPA predicted prominent activation of “pathogen induced cytokine storm signaling pathway” in aortas treated with Inact E (Figure 3B) or Act E+BAPN (Figure 3D). Other canonical pathways related to activation of monocytes and Th1 T-cells were also activated in the aortic wall under either condition (Figure 3B and 3D). Potent mediators common to acute inflammatory response, such as IFNG, TNF, and IL1B, were predicted to be significant upstream regulators for the differential gene expression provoked by Inact E or Act E+BAPN. In support of the predictions, IFNG, TNF, and IL1B were significantly upregulated by Inact E (Figure 3A, red dots) as well as Act E+BAPN (Figure 3D, red dots).

**Figure 3.**
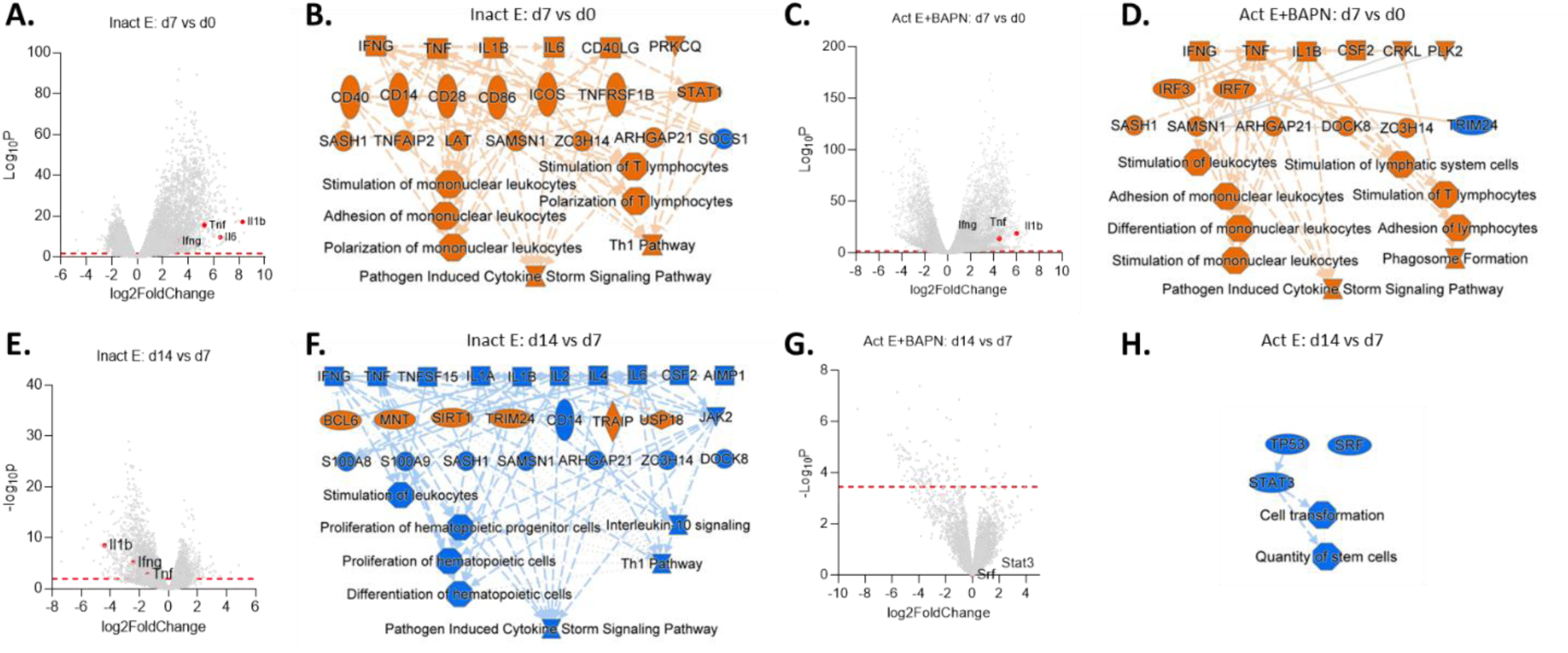
TADs failed to resolve the acute inflammatory response triggered by Act E+BAPN. A-D, genes (A and C) and pathways (B and D) induced by Inact E (A and B) and Act E+BAPN (C and D) at d7. Naïve ascending aortas served as the baseline reference. n=6/group. Volcano plots depict differentially expressed genes (DEGs) induced by Inact E (A) and Act E+BAPN (C). Potential upstream regulators and downstream pathways accountable for DEGs were identified using IPA and graphically summarized in B and D. Note the commonality of the downstream pathways activated by Inact E or Act E+BAPN, indicating prominence of a similar acute inflammation occurring under those conditions. E-H, temporal changes of gene expression (E and G) and downstream pathways (F and H) in aortas exposed to Inact E (E and F) or Act E+BAPN (G and H). Note the massive downregulation of genes from d7 to d14 in aortas exposed to Inact E only (E). Accordingly, all inflammatory pathways activated by d7 were inhibited in the following week (F), indicating resolution of acute inflammation. In contrast, only a few genes displayed a significant temporal change in expression in Act E+BAPN TADs (G), and these changes are attributable to an inhibition of SRF and TP53 (H)—regulators critical for SMC physiology, while changes in inflammatory pathways were not identified (H), indicating failure to resolve the acute inflammation. Dash red lines: levels of significant P; Brown: activation; Blue: inhibition; Red dot: actual expression of genes predicted to be significant upstream regulators by IPA.

Next, we investigated the temporal (i.e. d7 *vs.* d14) changes of gene expression in aortas challenged by Inact E or Act E+BAPN. 4828 genes displayed significant time-dependent expression in aortas exposed to Inact E (Figure 3E). Among these genes, 2714 genes were significantly downregulated (Figure 3E). In contrast, only 137 genes displayed significant temporal expression in aortas treated with Act E+BAPN (Figure 3G). IPA predicted an overwhelming inhibition of the inflammatory pathways activated at d7 for aortas exposed to Inact E (Figure 3F), while none of the inflammatory pathways activated at d7 in Act E+BAPN treated TADs was quenched by d14 (Figure 3H). SRF and TP53 were predicted to be inhibited (Figure 3H). However, these regulators are critical for SMCs to survive and maintain their contractile phenotype^29^. These results suggest that resolution of the acute inflammation might have contributed to the distinct phenotype observed for two groups at d28 (Figure 1).

### Both innate and adaptive immune responses are activated and intensified by Act E+BAPN in TADs

To determine specific molecular pathways responsible for TAD development induced by Act E+BAPN, we compared gene expression profile and analyzed DEGs between Act E+BAPN and Inact E aortas. At d7, 1457 DEGs were identified, with 801 and 656 genes being up- and down-regulated, respectively, in Act E+BAPN group compared with Inact E group (Figure 4A). IPA predicted activation of a set of upstream regulators to be accountable for the DEGs (Figure 4B). Most of them were expressed at a significantly higher level in Act E+BAPN TADs than in Inact E aortas (Figure 4A, red dots). It is worth of noting that IL17A and IL23A, both are critical players of IL17 signaling^30^, were predicted to be significant upstream regulators. TNF and IL1B, although both were upregulated by in aortas exposed to Inact E (Figure 3B) or in Act E+BAPN (Figure 3D), were upregulated to a significantly higher level in TADs compared with aortas treated with Inact E (Figure 4A, red dots). With this gene signature, IPA predicted activation of pathways specialized in “recruitment of myeloid cells”, including mononuclear cells, neutrophils, and phagocytes, in TADs (Figure 4B). By d14, 6390 DEGs were identified (Figure 4C), due to the resolution and the substantiation of the acute inflammation in Inact E and Act E+BAPN aortas, respectively (Figure 4C). Regulators of adaptive immunity were upregulated (Figure 4C, red dots) and predicted to promote Th1 (IFNG and IL12A) and Th2 (TSLP) responses (Figure 4D). Meanwhile, macrophages were regulated to differentiate and perform phagocytosis (Figure 4D), events commonly occur during chronic inflammation.

**Figure 4.**
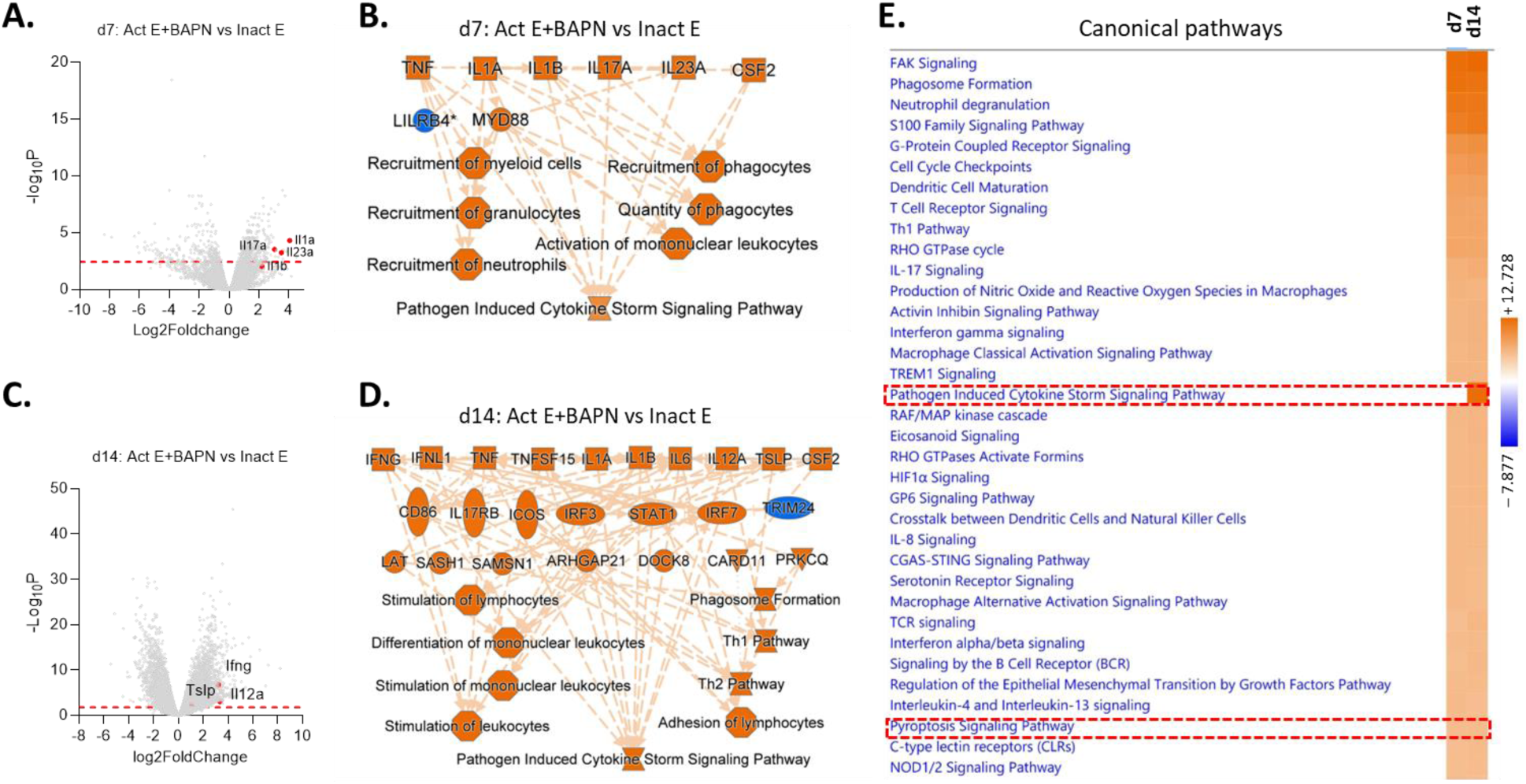
Act E+BAPN triggered a progressive immune response associated with enhanced pyroptosis signaling pathways in TADs. Comparisons were performed between Act E+BAPN and Inact E treatments—conditions both triggered an acute inflammation but resulted in distinct aortic phenotype. A-D, differential gene expression and pathway activation at d7 (A and B) and d14 (C and D). Expression of the predicted upstream regulators are highlighted in red in the volcano plots. Red lines: levels of significant P; Brown and blue: upstream regulators and pathways predicted to be activated and inhibited in Act E+BAPN TADs, respectively. Note the activation of neutrophil-mediated acute inflammation at the early stage (B) and T cell-mediated adaptive immune response at the relatively advanced stage (D). E, comparison of the relative activity of the significant pathways between d7 and d14. The heatmap covers only top pathways with a Z score ≥ 4.7. “Pyroptosis signaling pathway” emerged as one of the top-ranked cellular events (lower red dash line box).

Next, we analyzed enrichment of DEGs in KEGG pathways using DAVID functional annotation analysis. Because the maximum number of genes it takes is limited to 3,000, we only included DEGs with a fold change ≥ 2 and P ≤ 0.005. The enriched KEGG pathways were exported and plotted as dot plots (Figure 5A and 5B). Many of the enriched pathways were identified in IPA analysis (Figure 4E). However, additional signaling networks were uncovered. Among them, those related to stress response, cell growth, and cell differentiation, such as cAMP, cGMP-PKG, PI3K-AKT, MAPK, WNT, and regulation of actin cytoskeleton, were enriched on d7 (Figure 5A) and d14 (Figure 5B), indicating contribution of inflammation-independent mechanisms to TAD formation. This finding is particularly relevant to TAD formation since inflammation is not always present in human TADs^31^. Nevertheless, critical signaling for activation and differentiation of T- and B-cells, such as T- and B-cell receptor signaling, was not enriched at d7, but was at d14, suggesting activation of adaptive immunity in TADs. This finding is consistent with a previous study that CD4^+^ T-cells of smokers with TADs are autoimmune to elastin fragments^32^.

**Figure 5.**
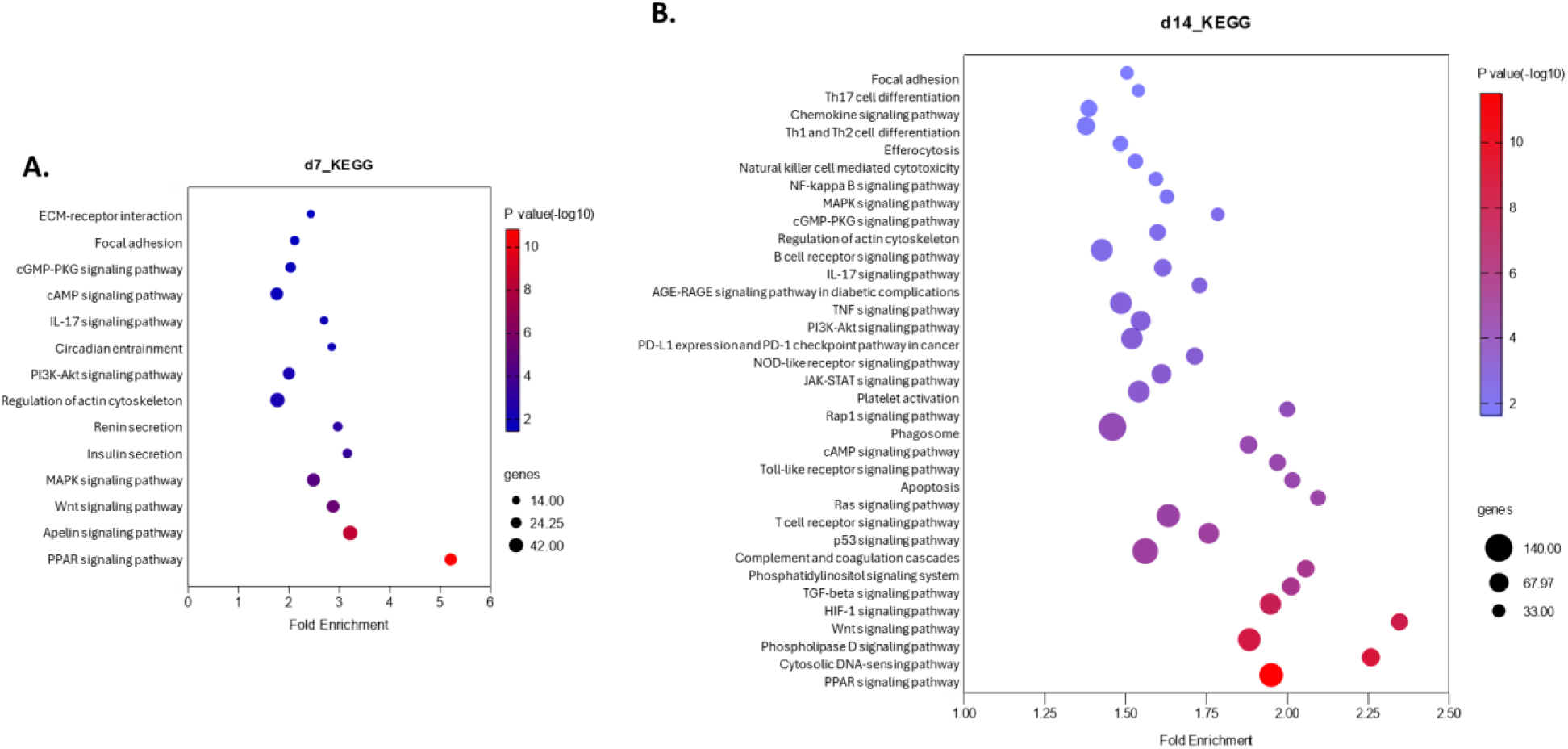
KEGG pathway analysis identified key signaling networks enriched in developing TADs. DEGs obtained with d7 and d14 TADs were analyzed with DAVID to discern KEGG signaling networks associated with activation of canonical pathways predicted by IPA. A, KEGG pathways enriched for d7 TADs. In addition to pathways that are generally considered to be proinflammatory, signaling networks related to stress response, cell growth, and differentiation, such as cGMP-PKG, cAMP, PI3K-AKT, MAPK, and WNT, were enriched in d7 TADs. B, KEGG pathways enriched for d14 TADs. While supporting activation of T-cell mediated adaptive immunity as predicted by IPA, the result demonstrated enrichment of the same set of “non-inflammatory” pathways as identified at d7, suggesting persistent activation of these pathways in TADs.

Finally, we performed IPA to assess the relative activity of the pathways in TADs between d7 and d14. A total of 288 canonical pathways had a significant Z score (i.e. Z ≥ 2.0), of which only 8 pathways were inhibited (Supplementary Table 1). Among the activated pathways, most of them are related to activation and differentiation of innate and adaptive immune cells, and “Pyroptosis Signaling Pathway” was ranked among the top 10 pathways specializing in regulation of cellular events and substantiated from d7 to d14 (Figure 4E). Given the proinflammatory nature of pyroptosis, it might play an important role in TAD development. Several studies have evaluated its role in AAA development^15,16,33^, but its effect on TAD formation had not been evaluated. Therefore, we addressed this issue.

### Cell death is an active event in developing TADs

TUNEL (terminal deoxynucleotidyl transferase dUTP nick end labeling) assays were performed to label dying cells in aortas exposed to different conditions for four weeks. TUNEL positive cells (TUNEL^+^) were located and distributed across all layers of the aortic wall in aortas exposed to Act E+BAPN, Act E, or Inact E +BAPN. While there appeared to be more TUNEL^+^ cells in the adventitial layer, a spatial pattern could not be established for any of the conditions due to the small number of TUNEL^+^ (Figure 6A). However, a significantly greater number of TUNEL^+^ cells were detected in aortas challenged with Act E+BAPN than those subjected to Act E, or Inact E+BAPN, while hardly any were noted in aortas treated with Inact E (Figure 6B). This differential degree of cell death correlated with the histological finding that aortas exposed to Inact E only were free of intimomedial tears (Figure 1C).

**Figure 6.**
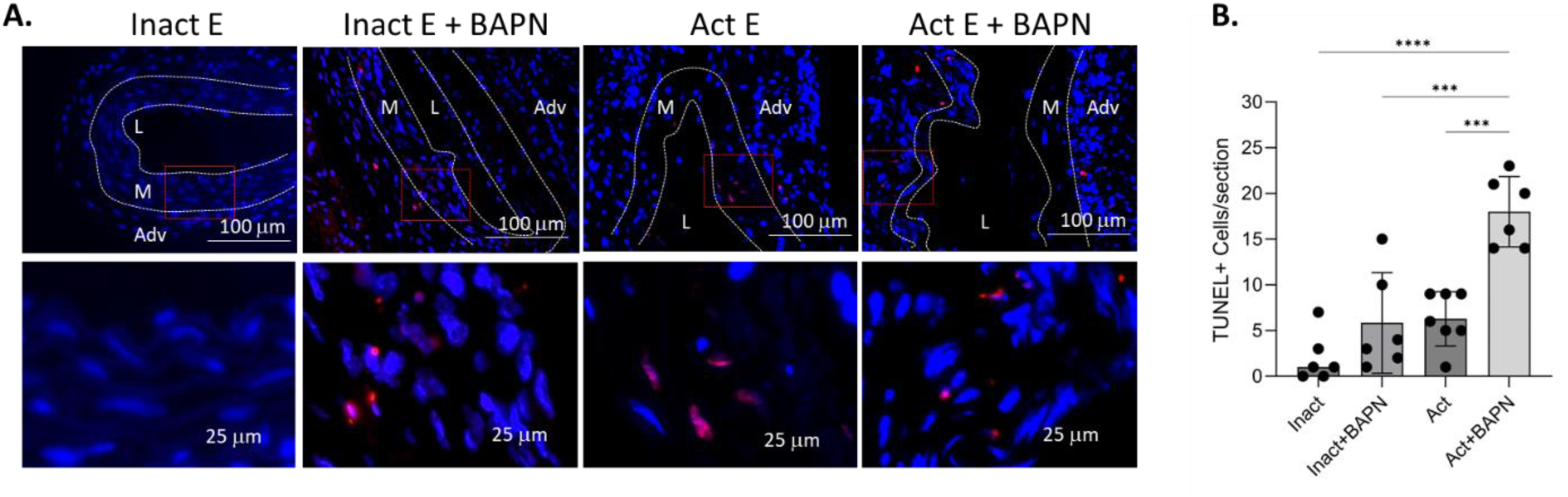
Act E+BAPN caused a greater degree of cell death than other conditions in d28 TADs. A, representative images of TUNEL staining of TADs induced by the indicated challenges. A magnified view of the boxed area in A is shown on the second row. Blue: DAPI nuclear counterstain; red: positive TUNEL staining; purple: TUNEL^+^ nuclei. L: lumen; M: media; Adv: adventitia. White dash lines: internal and external elastic laminae. B, counts of TUNEL^+^ nuclei. Data were analyzed using one-way ANOVA. ****P<0.001; ***P=0.001.

### Cell death mediated by GSDMD contributes to TAD development

Pyroptosis is executed by multiple members of the gasdermin family^13^. We focused our study on GSDMD as suggested by studies of AAA development. TADs were created in WT and Gsdmd null (Gsdmd^−/−^) mice and followed for four-weeks. During the follow-up, WT TADs exhibited a significantly slower rate of dilation than Gsdmd^−/−^ TADs, while TADs of either genotype displayed a significant progressive enlargement over time. By d28, WT and Gsdmd−/− TADs increased their lumen diameter by 31% and 68%, respectively (Figure 7A). Morphometric evaluation revealed significantly less medial loss and adventitial fibrosis for Gsdmd^−/−^ TADs compared with the WT controls (Figure 7B and 7C). Consistent with the attenuation of the aortopathy, fewer TUNEL^+^ cells (Figure 7D), CD4+ (Figure 7E), and CD8+ (Figure 7F) T-cells were detected in Gsdmd^−/−^ than in WT TADs, whereas differences CD19+ B-cells and CD68+ macrophages were insignificant between WT and Gsdmd−/− TADs. These results suggest that GSDMD-mediated cell death is a significant driver of TAD development and the protective effect of GSDMD deficiency likely results from modulation of a T-cell mediated immune response. This conclusion is also supported by the IPA prediction that Th1 and Th2 pathways are activated in TADs (Figure 4D).

**Figure 7.**
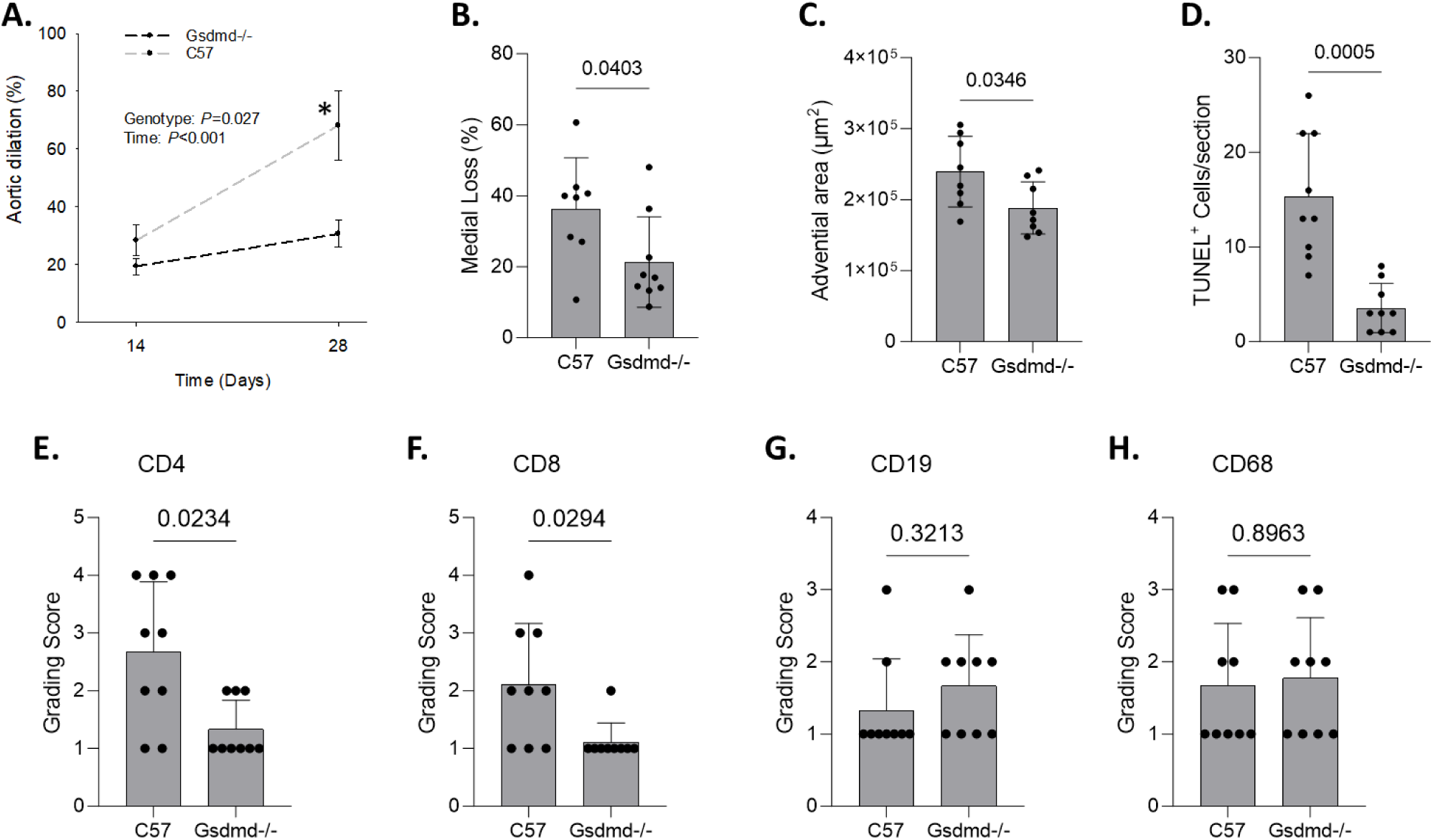
Genetic deficiency of GSDMD attenuated TAD formation. A: aortic dilation of TADs created with WT (C57) and Gsdmd null (*Gsdmd^−/−^*) mice (male, 10 to 14 weeks, n=15 in each group). Data were expressed as percent increase in lumen diameter from the baseline (i.e. pre-TAD induction) and analyzed using two-Way Repeated measurement ANOVA. *P=0.028, *Gsdmd^−/−^* vs. C57 at d28. B-C, morphometric evaluation with indicated parameters. D, semi-quantification of TUNEL+ cells. E-H, semi quantification of immunohistochemistry assays for cells with the indicated markers. Data were analyzed using unpaired t-tests.

### Inflammatory cell death programs are activated in human TADs

Previous studies from other laboratories demonstrated activation of NLRP3 inflammasome and caspase-1 in human TADs^34^. GSDMD is a substrate of Caspase-1. When cleaved, it releases pore-forming N-terminal fragments, causing cell death^13^. With an antibody specifically against the N-terminal fragment, cleaved GSDMDs were detected in the medial layer of TADs (Figure 8), indicating that GSDMD-mediated cell death programs are activated in human TADs.

**Figure 8.**
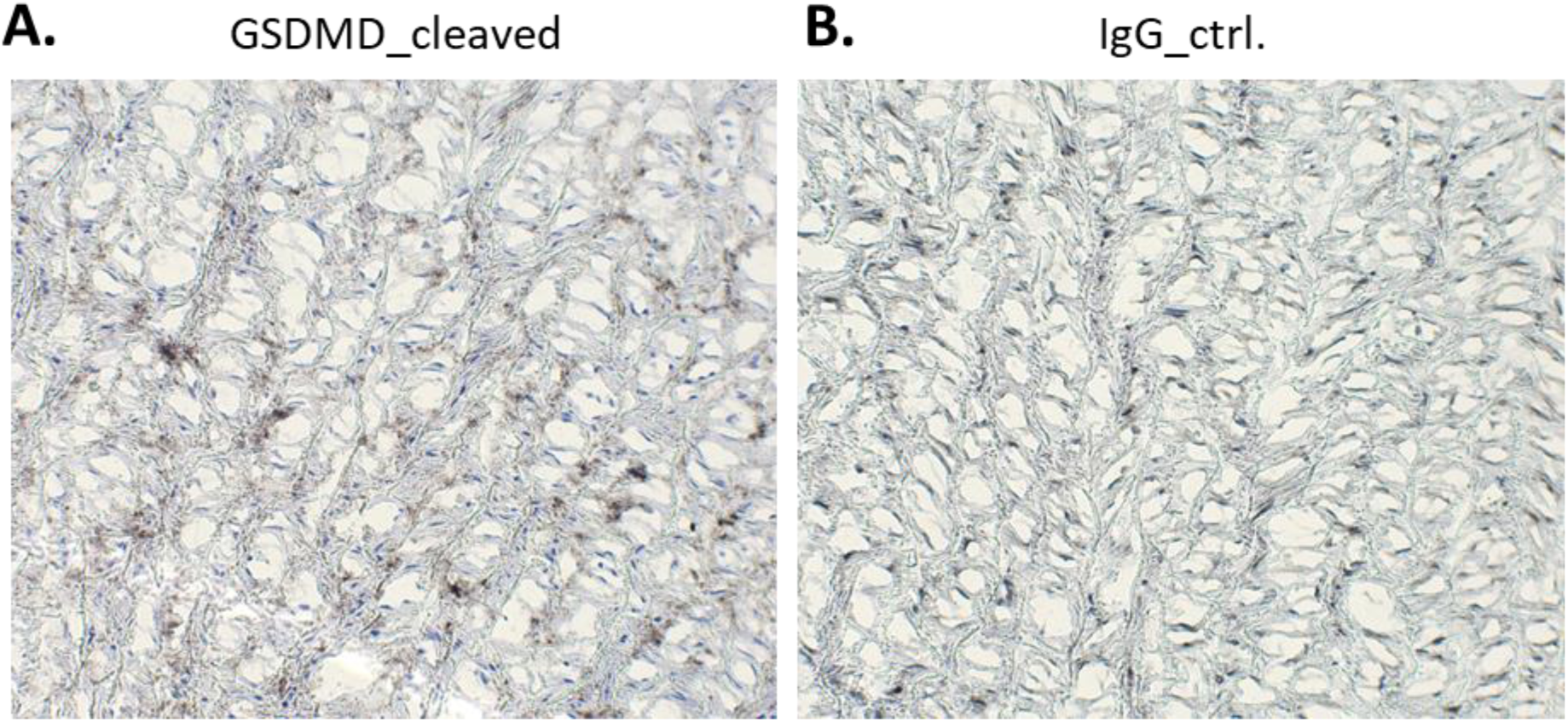
Cleaved GSDMDs and activation of multiple upstream pathways were detected in human TADs. A, IHC assays with a specific rabbit antibody against N-terminal fragments of cleaved GSDMDs. Brown, positive staining; Blue: nuclear counterstain; B, negative control with non-immunized rabbit IgGs.

## Discussion

In the present study, the model created recapitulates several critical pathologies of human TADs. It features an early (< 1 week) onset of intimomedial tears and dissection, followed by medial loss and adventitial fibrosis. Mechanistically, myeloid cells particularly neutrophils are likely the driver of the early pathology, while medial loss and progressive dilation during the late stage is likely to be perpetuated by chronic inflammation involving activation of both innate and adaptive immune responses. Enrichment of signaling networks related to cell growth, differentiation, and death, such as cAMP, cGMP-PKG, PI3K-AKT, GSDMD, is associated with these events. In support of these mechanisms, we demonstrated that GSDMD deficiency attenuated medial loss, adventitial fibrosis, and dilation of TADs. This attenuation is associated with reduced cell death and T-cell accumulation.

Because of the undefined etiology for the development of sporadic TADs, artificial challenges remain the only option for modeling the pathogenesis of TADs. Topical application of elastase in the descending aorta of mice induces TAAs when BAPN is not included in the induction^35,36^. A similar aortic phenotype can also be induced with topical application of calcium chloride^37^. However, these models only recapitulate pathologies of TAAs and do not develop phenotypic traits typical for aortic dissections. The widely adopted AngII model develops TADs^38^. We have reported that BAPN can be introduced to the system to enhance the rate of aortic rupture to 50% in WT mice^39^. One of the major limitations of this model is that it develops multiple isolated dissecting aneurysms with the location varying from the ascending to the abdominal aorta. Several studies^34,40,41^ including those from our laboratory^5,42^ showed that TADs and AAAs develop in the same animal and rupture at a similar rate in this model, which differs from the phenotype of human TADs that affect only the ascending (Type A) or spare the ascending (Type B) aorta^2^. TADs may be induced with BAPN. However, this model works only with immature animals^43–45^, which differs from the epidemiology of human TADs that affect most frequently the aging subpopulation, particularly those older than 60 years^2^. Our laboratory and other groups also demonstrated that SMC specific deletion of Tgfbr2^7^ or Tgfbr1^8,9,46^ spontaneously induces TADs in the ascending aorta. Major limitations of these models include time-consuming breeding processes to produce animals with compounded genetic modification (i.e. the Cre/loxP system) and off-target effect on expression of a large panel of genes at the baseline condition^10,47^. The TAD model created in the present study has overcome the limitations outlined above. It develops TADs at complete penetration. Additionally, the minimally invasive techniques applied in the model creation have successfully eliminated complications and mortality associated with the open-chest surgeries as previously documented in other studies^48^.

Quantification of the severity of TADs remains a challenge for studies using mouse models. For the AngII model, rate of rupture, incidence of dissection, aortic dilation, and complication of TADs have been taken into the equation^34,38,49^. This set of parameters represents the gold-standard for estimation of severity of TADs in the AngII model^40^. When using this model, we noticed that one of the hall-mark pathologies of the TADs is the partial or complete loss of the medial layer^5^. While this is not a typical pathology of human TADs, it is a factor to be considered for evaluation of medial degeneration. In previous studies, we used a five-scale scoring system, including medial loss, to evaluate severity of TADs induced by SMC-specific deletion of *Tgfbr1*^8^ or AngII+BAPN^5^. Significant medial loss in TADs have also been documented for experimental TADs by others^7,50^. In the present study, we showed that medial loss is a useful parameter for quantification of TADs induced by Act E+BAPN.

To meet the challenge of evaluating TAD severity, several studies have demonstrated advantages of using sophisticated imaging to detect and quantify the onset and progress of aortic dissection. These studies demonstrated the robustness of their imaging systems to detect pathologies that are not easily caught in the routine histological evaluation and to quantify the progressive spreading of dissections^51,52^. However, this approach is not readily adopted by other laboratories due to the requirement of special imaging expertise and sophisticated imaging instruments. A simple, sensitive, reproducible, and clinically meaningful system remains to be developed for evaluation of TAD severity in experimental studies.

Immune injury appears to be the primary driver of TAD formation induced by Act E+BAPN. The importance of inflammation in TAD formation has been widely accepted^53^. However, many of the studies focused on general inflammation, characterized by elevated levels of inflammatory cytokines, proteases, and macrophage accumulation. A few studies explored the effect of immune cells, such various subsets of T-cells, B-cells, and macrophages in the development of abdominal aortic aneurysms (AAAs) and came up with a hypothesis that Th2 and Th17 immunity is deleterious to AAA development^54,55^. In contrast, Th1 response is associated with TAAs^56–58^ and TADs^32,59^. In the present study, we showed that general inflammation with enhanced expression of TNFα, IL1β, IL6, and IFNG does not necessarily cause aortopathy. It is a unique type of inflammatory response specializing in activation and recruitment of neutrophils that correlated with the onset of TADs. This finding is consistent with the study by Kurihara et al, showing neutrophil MMP9 promoted acute aortic dissections of the thoracic aorta pre-treated with BAPN^43^.

Innate immune response triggered by DAMPs (damage associated molecular patterns) contributes to TAD formation. DAMPs are released by stressed or dying cells and serve as alarmins to activate protective mechanisms to maintain tissue homeostasis. However, the danger signals can be overshot and become pathogenic when uncontrolled^60^. Previous studies from other laboratories have demonstrated a critical role for self-DNA^49^ and S100A12^61,62^ in TAD formation. In the present study, we demonstrated that danger signals and pyroptosis pathways were activated at the early stage (≤ 1 week) in both TADs (i.e. challenged with Act E+BAPN) and aortas with intact structures (i.e. exposed to Inact E), indicating that these events alone are not sufficient to induce TAD formation. At the advanced stage, inflammation in aortas exposed to Inact E was nearly completely resolved, while substantiated with intensified pyroptosis pathways and activation of Th1 as well as Th2 immune responses in TADs. This perpetuation of the inflammatory response correlated with a progressive medial degeneration and aortic dilation, suggesting that pyroptosis may play a role in subsequent remodeling, rather than initial onset of TAD formation.

Pyroptosis is executed by GSDMs of which the cleaved N-terminal fragments form pours in the cell membrane, leading to membrane rupture and cell death^13^. Using the TAD model created in the present study, we demonstrated that genetic deficiency of GSDMD attenuated TAD development by slowing medial loss and aortic dilation at an advanced stage. The attenuation is associated with mitigation of T-cell accumulation in TADs, suggesting a role for GSDMD-mediated inflammatory cell death in modulating T-cell response and TAD remodeling. Studies of other laboratories have demonstrated protective effect of inhibition of GSDMD on AAA formation via pyroptosis-dependent^16^ and -independent^15^ mechanisms. Whether these mechanisms apply to TAD formation remains to be determined. It is also note that GSDMD is also an executioner of panoptosis and can be activated by multiple panoptosomes^14^. Future studies are warranted to evaluate activation of this pathway in TADs.

A few limitations exist in the present study. First, TADs induced by Act E+BAPN rupture at a very low rate in four weeks, and therefore, may not be applicable to studies focusing on aortic rupture. Although dissections occurred in >70% (7 of 9) TADs in the first week, they tend to advance to a chronic medial destruction, a phenotype like unruptured TADs induced by AngII. Furthermore, our model appears to conflict with previous reports that Act E with or without BAPN causes formation of aortic aneurysms in abdominal^27,28^ or descending thoracic aorta^26^. This discrepancy might result from anatomic difference of aortas in response to exogenous stimulation, which is supported by a very recent study demonstrating that BAPN causes distinct pathology in ascending and descending aortas^45^. Next, our results suggest that a unique type of inflammatory response, particularly the recruitment of neutrophils, drives the onset of TAD formation. While this finding is consistent with results obtained by others using the AngII model^43^, whether it is MMP9 and/or other mechanisms that cause the early dissection remains to be addressed. Additionally, whether medial loss results from dissection (i.e. physical removal of the medial layer), chronic death of SMCs, and/or other events remains to be determined. Finally, GSDMD may be activated via the inflammasome^13^ and panoptosome pathways^63^. Further studies are needed to determine the upstream pathways that lead to cleavage of GSDMD in TADs.

## Acknowledgments

None

